# Membrane insertion of soluble CLIC1 into active chloride channels is triggered by specific divalent cations

**DOI:** 10.1101/638080

**Authors:** Lorena Varela, Alex C. Hendry, Encarnacion Medina-Carmona, Diego Cantoni, Jose L. Ortega-Roldan

## Abstract

The CLIC family of proteins display the unique feature of altering their structure from a soluble form to a membrane-bound chloride channel. CLIC1, a member of this family, can be found in the cytoplasm or in nuclear, ER and plasma membranes, with membrane overexpression linked to tumour proliferation. The molecular switch promoting CLIC1 membrane insertion has been related to environmental factors, but still remains unclear. Here, we use solution NMR studies to confirm that both the soluble and membrane bound forms are in the same oxidation state. Our data from fluorescence assays and chloride efflux assays indicate that Ca^2+^ and Zn^2+^ trigger association to the membrane into active chloride channels. We use fluorescence microscopy to confirm that an increase of the intracellular Ca^2+^ leads to re-localisation of CLIC1 to both plasma and internal membranes. Finally, we show that soluble CLIC1 adopts an equilibrium of oligomeric species, and Ca^2+^/Zn^2+^ mediated membrane insertion promotes the formation of a tetrameric assembly. Thus, our results identify Ca^2+^ and Zn^2+^ binding as the molecular switch promoting CLIC1 membrane insertion.

**SIGNIFICANCE STATEMENT:** CLIC1, a member of the CLIC family of proteins, is expressed as a soluble protein in cells but can insert in the membrane forming a chloride channel. This chloride channel form is upregulated in different types of cancers including glioblastoma and promote tumour invasiveness and metastasis. The factors promoting CLIC1 membrane insertion nor the mechanism of this process are yet understood. Here, we use a combination of solution NMR, biophysics and fluorescence microscopy to identify Ca^2+^ and Zn^2+^ binding as the switch to promote CLIC1 insertion into the membrane to form active chloride channels. We also provide a simple mechanism how such transition to the membrane occurs. Such understanding will enable subsequent studies on the structure of the chloride channel form and its inhibition.

## INTRODUCTION

The Chloride Intracellular Channel (CLIC) family consists of a group of highly homologous human proteins with a striking feature, their ability to change their structure upon activation from a soluble form into a membrane bound chloride channel, translocating from the cytoplasm to intracellular membranes (1, 2). CLIC1 is the best characterised of the CLIC protein family. It is expressed intracellularly in a variety of cell types, being especially abundant in heart and skeletal muscle (2). CLIC1’s integral membrane form has been found to be localised mostly in the nuclear membrane, although it is present in the membranes of other organelles and transiently in the plasma membrane. It has also been shown to function as an active chloride channel in phospholipid vesicles when expressed and purified from bacteria, showing clear single channel properties (3, 4).

CLIC1 has been implicated in the regulation of cell volume, electrical excitability (5), differentiation (6), cell cycle (7) and cell growth and proliferation (8). High CLIC1 expression has been reported in a range of malignant tumours, including prostate (9), gastric (10), lung (6) and liver (11) cancers, with evidence of CLIC1 promoting the spread and growth of glioblastoma cancer stem/progenitor cells (12, 13).

The activity and oncogenic function of CLIC1 is modulated by its equilibrium between the soluble cytosolic form and its membrane bound form. Only CLIC1 in its channel form has been shown to have oncogenic activity, and specific inhibition of the CLIC1 channel halts tumour progression (13). However, to date very little and conflicting information is available for the membrane insertion mechanism, and the structure of the channel form is unknown. Oxidation with hydrogen peroxide causes a conformational change due to the formation of a disulphide bond between Cys24 and the non-conserved Cys59, exposing a hydrophobic patch that promotes the formation of a dimer (14), in a process that has been proposed to lead to membrane insertion (15). However, numerous studies have shown that oxidation does not promote membrane insertion(16); with evidence pointing at pH (17, 18) or cholesterol (19) as the likely activation factors. Thus, long standing inconsistencies in the data surrounding the molecular switch that unusually transforms CLIC1 from its soluble form into a membrane bound channel has prevented further advances in the understanding of CLIC1 function. In this study, we have explored the membrane insertion mechanism of CLIC1. We used NMR experiments on CLIC1 extracted from the soluble and membrane fraction of *E. coli* to show that membrane bound CLIC1 is in a reduced state. We demonstrate that CLIC1 exists in an equilibrium between monomers, dimers and higher-order oligomers, and that oligomerisation of CLIC1 is required for membrane insertion. Finally, we use membrane binding assays to show that divalent cations enhance CLIC1 membrane insertion, and fluorescence microscopy and chloride efflux assays to confirm that 2+ cation-triggered membrane insertion results in the formation of active chloride channels.

## RESULTS

### Membrane insertion is not driven by oxidation

CLIC1 has previously been successfully expressed in *E. coli*, purified and assayed for chloride conductance (1). To assess if recombinant CLIC1 is able to insert in *E. coli* membranes, we expressed a C-terminal GFP tagged construct in *E. coli* and isolated both the cytosolic and membrane fractions. GFP fluorescence measurements for both the soluble fraction and membranes resuspended in similar volumes indicate that the majority of recombinant CLIC1 inserts in the *E. coli* membrane (Figure 1A). CLIC1 is a human protein, but it has been observed to possess chloride efflux activity in mixtures of lipids containing Phosphatidyl Serine (PS) (4). Given the high proportion of PS lipids in *E. coli* membranes, it is not surprising that CLIC1 can insert into bacterial membranes.

**Figure 1.**
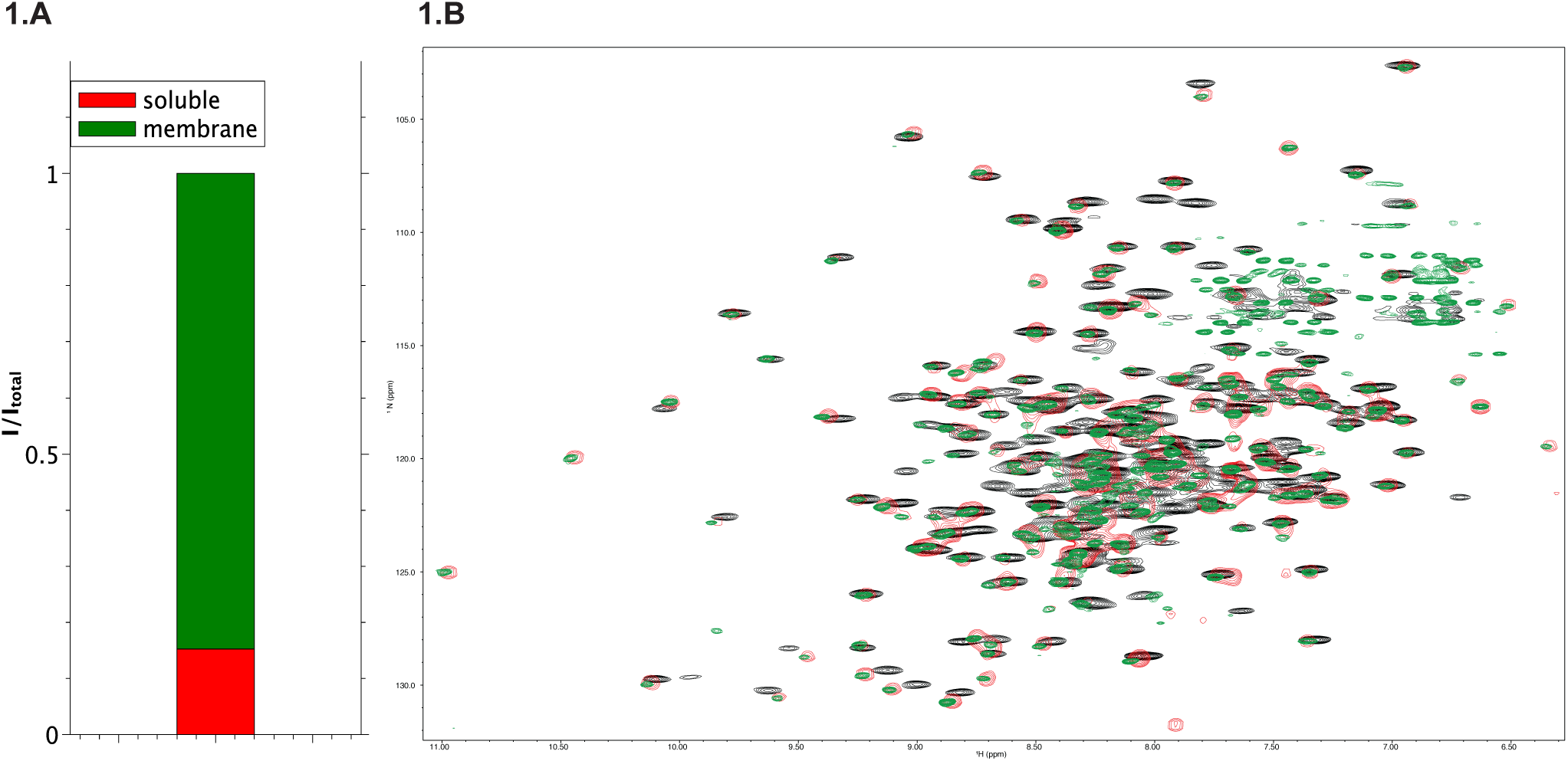
CLIC1 membrane insertion in *E. coli* is not driven by oxidation. (A) - Quantification of CLIC1 extracted from the membrane (green) or the soluble fraction (red). (B) – Overlay of ^15^N Trosy HSQC or SoFAST HMQC spectra of CLIC1 extracted and purified from the *E. coli* membrane fraction (green), or cytosol (red) and oxidised with 50 mM H_2_O_2_ (black).

Oxidation has been proposed as the key trigger for membrane insertion, through the formation of a disulphide bridge between the conserved Cysteine 24 to the non-conserved Cysteine 59, although more recent studies do not reconcile with this mechanism (16, 19). To test the oxidation state of both soluble and membrane fractions, ^15^N-labelled CLIC1 was expressed recombinantly in *E. coli* and purified from the membrane and soluble fractions independently. 2D ^15^N TROSY or ^15^N-SOFAST-HMQC experiments were collected for each fraction. ^1^H and ^15^N chemical shifts were measured, as the chemical shifts of NH moieties are very sensitive to dynamics, as well as the local and global structure of the protein. Any structural changes resulting from disulphide bond formation would have a big impact in the chemical shifts of a large subset of NH resonances. An overlay of spectra from both fractions shows that the CLIC1 proteins they contain are nearly indistinguishable (Figure 1B), indicating that both forms are in the same oxidation state. Reduction of both samples with 5 mM DTT did not cause significant alterations in the spectrum (Figure S1). In contrast, oxidation with H2O_2_ resulted in large chemical shift differences for a subset of resonances in both spectra, indicating that CLIC1 is inserted in *E. coli* membranes in the reduced state, and therefore the membrane association process is not triggered by oxidation.

### Divalent cations trigger CLIC1 membrane insertion

Since our data indicates that oxidation does not induce membrane insertion, we screened for different conditions that could trigger membrane insertion. A membrane insertion assay was developed, in which CLIC1 was mixed with the lipid mixture asolectin. The mixture was subsequently ultra-centrifuged to separate the soluble and membrane bound components. Native tryptophan fluorescence experiments were collected from the initial mixture, the supernatant and membrane pellets fractions. Using phosphate buffer, which forms insoluble complexes with divalent cations, no insertion could be detected even in the presence of oxidizing conditions. (Figure 2A). A simple change in the buffer composition to HEPES resulted in an increase in the tryptophan fluorescence emission spectrum for the membrane fraction. Since phosphate is known to form insoluble complexes with divalent cations, we explored whether the lipid insertion could be triggered by binding of divalent cations. A series of membrane insertion assays were conducted in the presence of Zn^2+^, Ca^2+^ and Mg^2+^ to identify the effect of 2+ metals (Figure 2A). A significant increase in the overall intensity in the emission fluorescence spectra of the membrane fractions of CLIC1 was found in samples incubated with Zn^2+^, and in a lower extent with Ca^2+^, suggesting that CLIC1 membrane insertion is driven by binding to Zn^2+^ and/or Ca^2+^. In a further series of experiments, tryptophan emission fluorescence spectra were measured at increasing concentrations of Zn^2+^ in the presence and absence of asolectin vesicles. In the absence of lipids, a decrease in the overall intensity of fluorescence is observed, consistent with aggregation of the protein that eventually lead to the appearance of a precipitate. In the presence of lipid vesicles, a blue shift of the fluorescence maximum is observed which was consistent with a lack of aggregation, confirming that divalent cations trigger membrane insertion or association and ruling out any interference of protein aggregation in our membrane insertion assay (Figure S2). To confirm this interaction with lipid bilayers, fluorescence microscopy images were taken in mixtures of GFP-labelled CLIC1 and giant unilamellar vesicles (GUVs) labelled with Nile red dye. While in the absence of divalent cations no co-localisation of CLIC1 and vesicles was found (Figure 2C), addition of Zn^2+^ resulted in complete co-localisation of CLIC1 and the GUVs (Figure 2C and Figure S3A,B). Ca^2+^ ions also promote CLIC1 membrane association, with a lower level of co-localisation (Figure 2C and Figure S3C,D).

**Figure 2.**
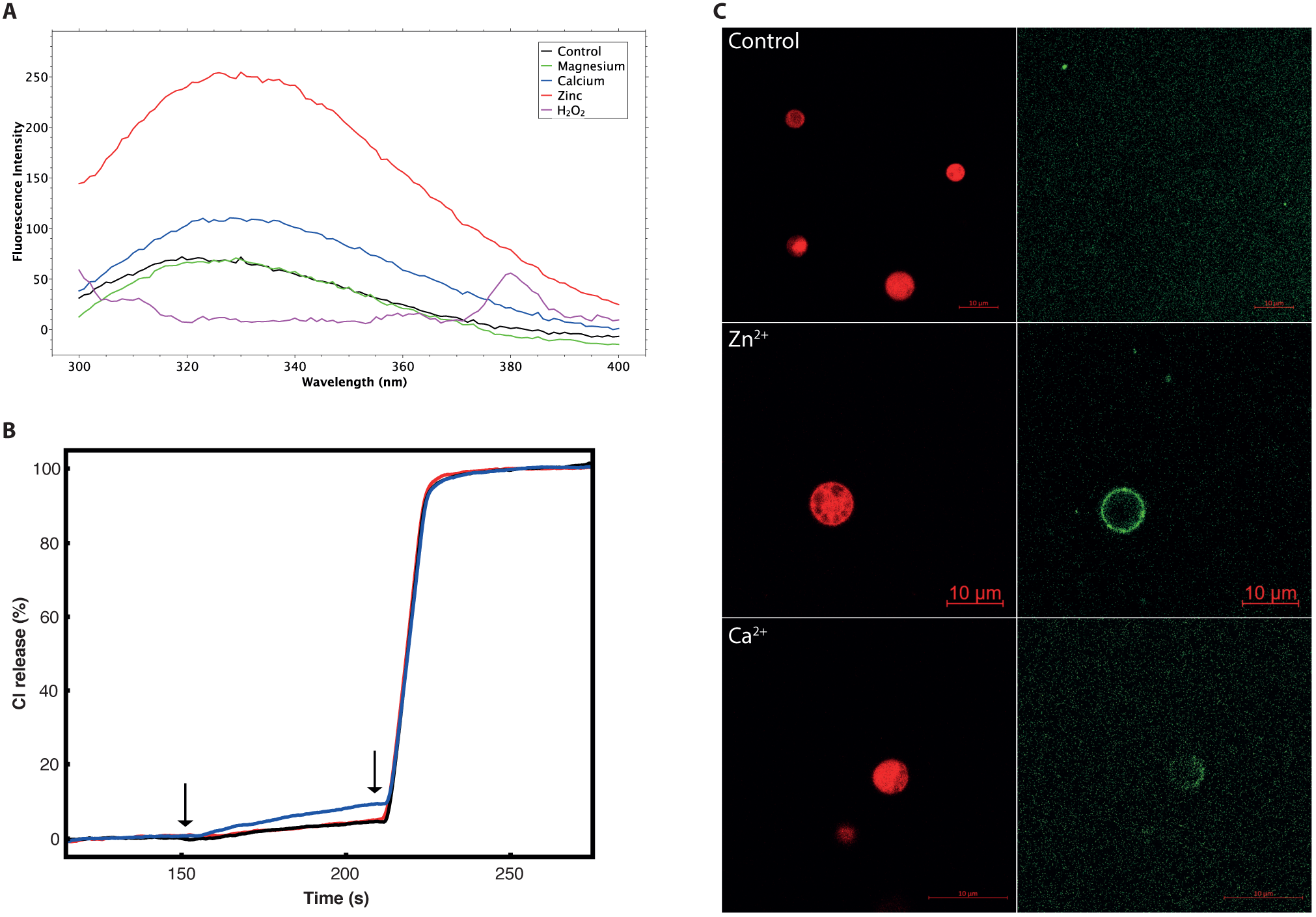
Divalent cations trigger CLIC1 membrane insertion. A - Native tryptophan fluorescence emission spectra of the membrane fraction of asolectin samples incubated with CLIC1 in the presence of H_2_O_2_ (magenta), no divalent cations (black), Ca^2+^ (blue), Zn^2+^ (red) and Mg^2+^ (green). B -CLIC1 chloride conductance monitored in Asolectin vesicles in presence of Zn^2+^ (blue) or EDTA (red). A control experiment without the addition of CLIC1 is represented in black. The first arrow indicate the addition of Valinomycin, and the second arrow indicates the addition of Triton X-100. C – Fluorescent microscopy images of Asolectin GUVs labelled with Nile red dye incubated with GFP-labelled CLIC1 in the absence of metals (top) and in the presence of 500 μM of Zn^2+^ (middle) or Ca^2+^ (bottom). The left panel shows images exciting Nile red, and the left panel shows images exciting GFP.

To test if membrane bound CLIC1 possess chloride transport properties, chloride efflux was recorded upon addition of Valinomycin using CLIC reconstituted in asolectin vesicles in the presence of Zn^2+^ and Ca^2+^.. While CLIC1 shows chloride efflux activity in the presence of Zn^2+^ (Figure 2B) or Ca^2+^ (Figure S4), incubation with EDTA leads to the complete repression of chloride efflux, confirming that efflux observed with divalent cations is due to the formation of active CLIC1 channels.

We sought to examine the influence of the increase of these metal cations on mammalian cell lines in the presence of CLIC1 and whether *in vivo*, CLIC1 would localise to the plasma or internal membranes. To investigate this, we stained endogenous CLIC1 in HeLa cells with CLIC1-specific antibodies, and treated with 5 mM extracellular Ca^2+^ or 10μM Ionomycin to increase the intracellular Ca^2+^ concentrations. The intracellular calcium levels were monitored using a Fluo4 reporter system. A moderate increase of intracellular Ca^2+^ levels could be observed upon stimulation with extracellular Ca^2+^, and a marked increase was found upon stimulation with Ionomycin (Figure S5). Fluorescence microscopy demonstrated that CLIC1 localisation changes in HeLa cells upon increasing intracellular calcium levels (Figure 4 and Figure S6). Control cells with no addition of Ca^2+^ show a cytoplasmic localisation of CLIC1 with little observable membrane localisation. Contrastingly, an increase in the intracellular Ca^2+^ levels results in CLIC1 re-localisation to internal membranes and the plasma membrane and a notable decrease of CLIC1 concentration in the cytoplasm, demonstrating the effect of 2+ metal cations on CLIC1 localisation in cells. Together with our previous findings, these images show that metal cations are imperative to CLIC1’s mechanism of insertion into the membrane and control of where it is localised.

**Figure 3.**
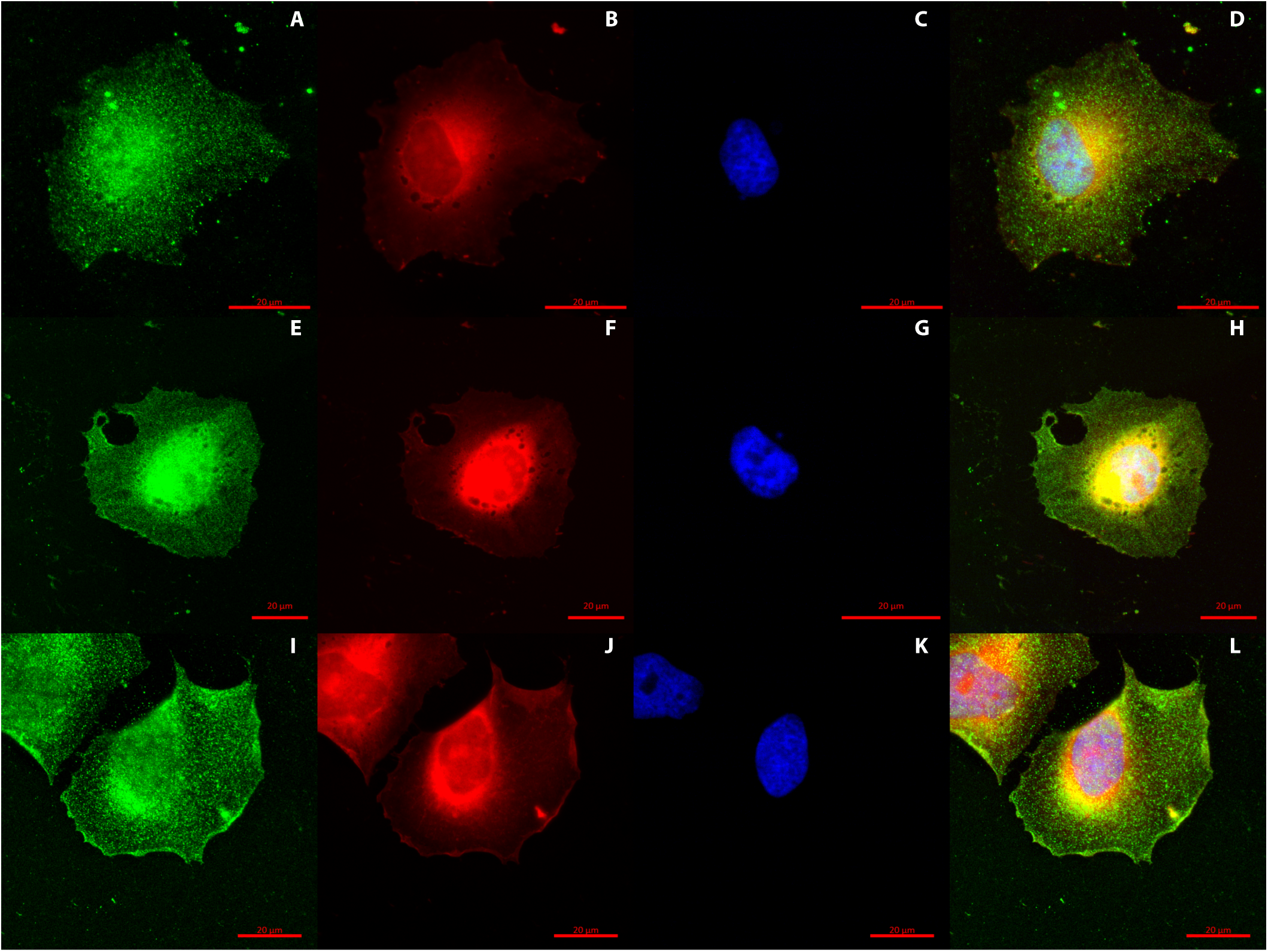
Ca^2+^ driven re-localisation of CLIC1 in HeLa cells. Fluorescence microscopy images of HeLa cells stained with a CLIC1 antibody (green), a membrane marker (red) and DAPI (blue) in the absence (A-D) and presence (E-H) of 5 mM Ca^2+^ in the media or upon treatment with 10 μM Ionomycin (I-L). Images in D, H and L show a merge of the three different channels for the three conditions.

**Figure 4.**
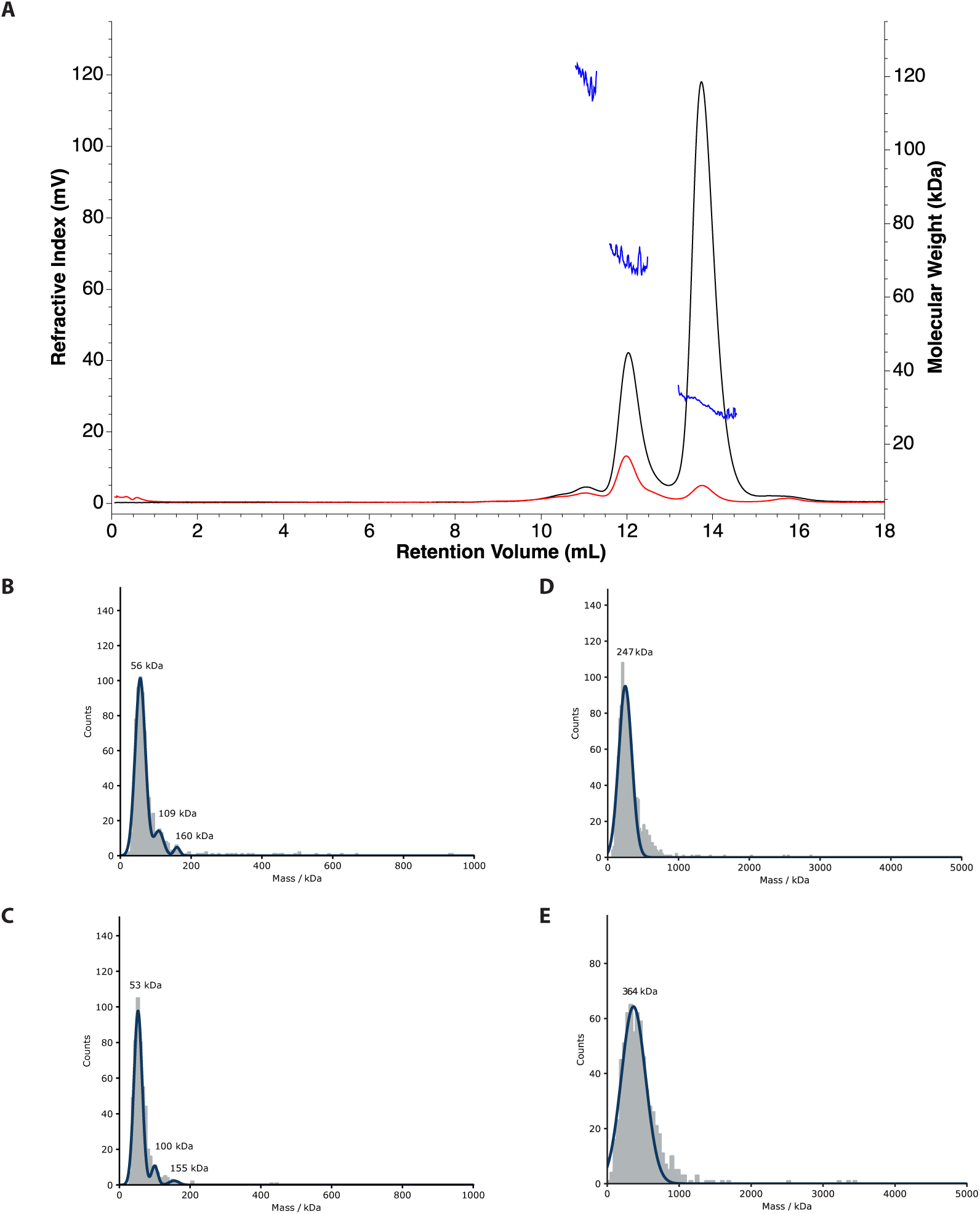
CLIC1 oligomeric states in solution and in the chloride channel form. SEC-MALS traces of non-treated CLIC1 (black). CLIC1 samples corresponding to the dimeric form were subjected for a second SEC-MALS run (red). The molecular weights calculated for the monomer, dimer and tetramer peaks are shown in blue. B,C – Mass Photometry histograms for 100nM CLIC1 samples in the absence and presence of an equimolecular concentration of Zn^2+^. D,E – Mass Photometry histograms of empty and CLIC1-containing nanodiscs.

### Dynamics of CLIC1 in solution shows oligomerization in equilibrium

The formation of the CLIC1 channel has been shown to involve oligomerisation in the membrane to large complexes containing six to eight subunits (20). In light of this, we explored if CLIC1 oligomerisation also occurs in solution, and if it was modulated by divalent cation binding. Size exclusion chromatography (SEC) in reducing and not-reducing conditions indicates the presence of at least three species with different molecular weight and some minor high-order oligomers that are not dependent on the formation of disulphide bonds (Figure 4 and S7). To assess the stability of the monomers and oligomers both species was then subjected to a second SEC step. Again, two peaks were obtained, indicating that CLIC1 exists in a non-covalent equilibrium between the two species (Figure 4A). This was confirmed by multiangle light scattering (MALS). SEC-MALS analysis indicates that the main peaks correspond to monomeric, dimeric and tetrameric species. Treatment with Zn^2+^ resulted in strong interactions with the Superdex200 matrix, and no elution peak could be detected.

CLIC1 samples were also subjected to interferometric scattering mass photometry (iSCAMS). Due to the limit of detection being in the range of 40KDa no monomers could be observed, and only dimers and tetramers and hexamers were detected (Figure 4B). Treatment with equimolar concentrations of Zn^2+^ did not significantly alter the equilibrium between oligomeric species in solution, ruling out that CLIC1 oligomerisation occurs as a consequence of divalent cation activation (Figure 4C). To study the oligomeric state of CLIC1 in the membrane bound form, soluble CLIC1 was incubated with asolectin vesicles in the presence of Zn^2+^, and the lipid fraction was treated with SMAs to form nano-discs. Mass photometry measurements with empty and CLIC1 containing discs indicated a shift in the average mass of around 100KDa, consistent with a tetrameric assembly in the chloride channel state in nano-discs (Figure 4D-E). A similar shift is observed upon incubation of empty nano-discs with 1μM Zn^2+^.

## DISCUSSION

### A model for CLIC1 membrane insertion

Combining our data we can propose a new mechanism of CLIC1 membrane insertion (Figure 5) whereby soluble CLIC1 exists as a mixture of oligomeric states, mainly monomers, dimers and tetramers, exploring the oligomeric state of the membrane-bound form. Upon intracellular Ca^2+^ release (or release of other divalent cations), CLIC1 alters it structure likely exposing a hydrophobic segment and inserts in the membrane in a tetrameric assembly, forming active chloride channels. The propensity to oligomerise in solution, exploring the same tetrameric association than in the membrane bound form, contributes to diminish the entropic penalty of this assembly. CLIC1 binding to divalent cations, on the other hand, does not contribute to the oligomerisation process, and likely contribute to the exposure of a hydrophobic region on the N-terminus of the protein. Previous work (20) using FRET and oxidation suggested an oligomeric model comprising hexamers or octamers. While the formation of SMALPs could impact the oligomeric state of the chloride channel, the direct detection of mass by interferometric scattering mass spectrometry on native, unoxidized CLIC1 prepared inserting CLIC1 in vesicles or directly in SMALPs give us confidence in the validity of our measurements

**Figure 5.**
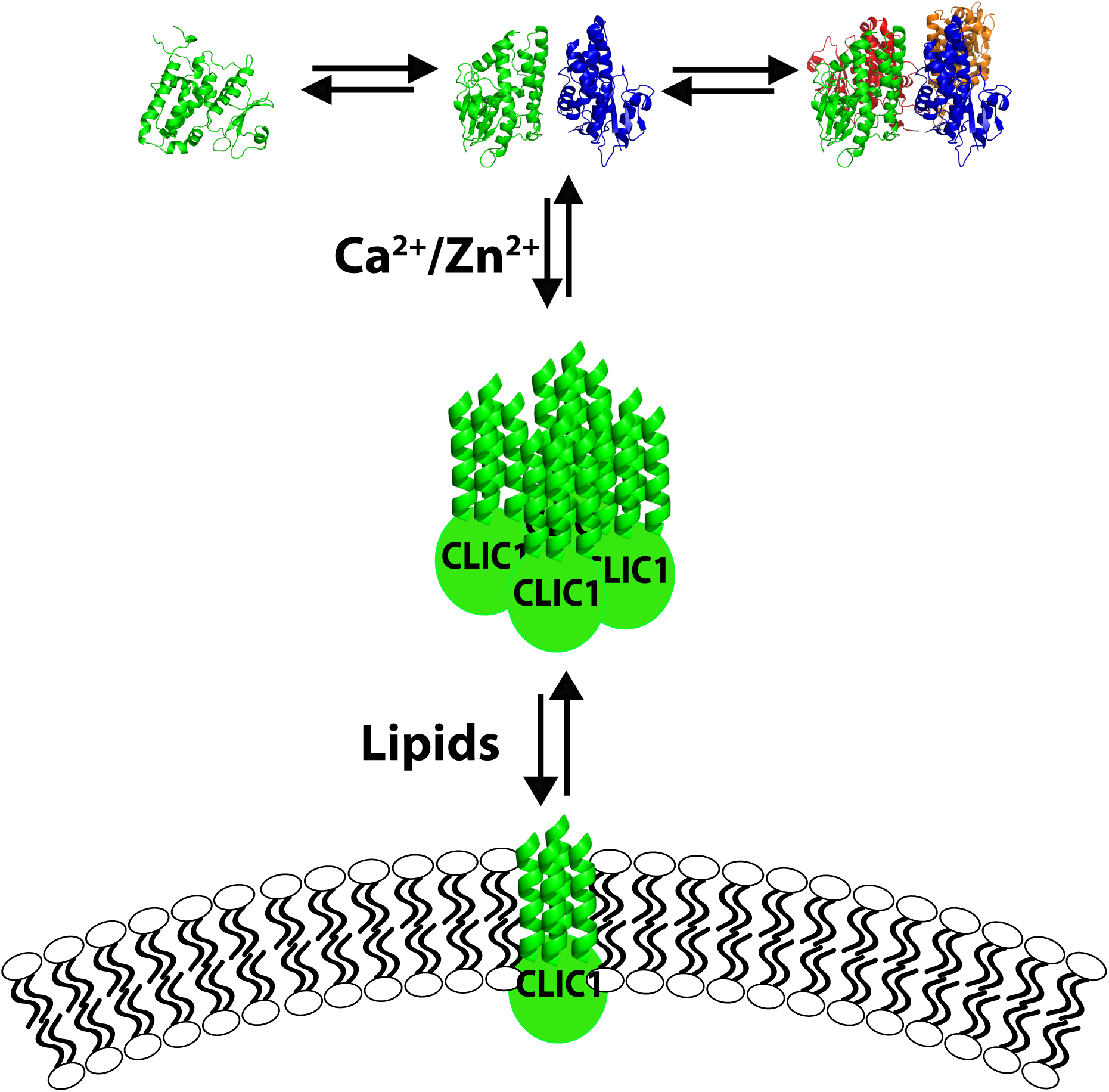
A model of the membrane insertion mechanism of CLIC1. Cytosolic CLIC1 exists in solution (Structural model from PDB 1k0m). With an increase in the intracellular Ca^2+^ (or Zn^2+^) levels, CLIC1 changes its structure, associates and insert in the membrane, forming active chloride channels.

While Zn^2+^ triggers a more pronounced insertion of CLIC1 in the membrane than Ca^2+^, a rise in intracellular Ca^2+^ is sufficient for CLIC1 re-localisation to the membrane. An increase in the concentration of both cations has been linked upstream and downstream of the ROS signalling pathway (21, 22), explaining why CLIC activity has previously been related to ROS production and oxidative stress. Further work is required to address this cation selectivity, but one could hypothesise that maximal membrane insertion of CLIC1 could be detrimental for the cells. Ca^2+^ on the other hand would enable a better regulated equilibrium between soluble and membrane bound CLIC1.

Calcium has been related to the membrane insertion properties of proteins of the annexin family (23), as well as the E1 membrane protein of rubella virus (24) and the amyloidogenic peptide amylin (25), promoting the interactions of the soluble forms of these proteins with negatively charged lipids. The extract of soy bean lipids asolectin, which is rich in the lipid classes PE, PC and the negatively charged PI, has been shown to promote maximal CLIC1 chloride efflux activity (4), supporting the role of divalent cations in CLIC1 membrane insertion. Annexin membrane association is expected to occur due to the exposure of an otherwise buried amphipathic segment upon binding to calcium ions. CLIC1 contains a region (residues 24-41) with moderate hydrophobicity and a moderate hydrophobic moment that could, in a similar mechanism, detach from the protein’s globular structure upon divalent cation binding, become exposed to the solvent and mediate association with the membranes, likely forming a helix. The hydrophobicity of this segment would also explain the aggregation of the protein at higher concentrations of calcium and zinc ions in the absence of lipids. Interestingly, the same region in the structurally homologous Glutathione S-transferase is not hydrophobic, suggesting that this helix plays a different role in the CLIC family.

While the structural rearrangements involved in this process are not yet fully understood, the molecular switch between the soluble and membrane bound forms and the changes in oligomerisation state required to generate the chloride channel assembly are now elucidated. This provides a clear mechanism for this unusual and clinically important channel formation process.

## METHODS

### Protein Expression and Purification

The Human CLIC1 gene (clone HsCD00338210 from the Plasmid service at HMS) was cloned into a pASG vector (IBA) containing an N-terminal twin strep tag and into a pWaldo (26) vector containing a C-terminal GFP. CLIC1 was expressed recombinantly in the C43 *E*.*coli* strain (Lucigen). The cells were lysed by sonication, and the membrane and soluble fractions were separated by ultracentrifugation at 117734 g. Membrane-bound CLIC1 can be extracted using a mixture of 1% DDM (Glycon) and 1% Triton X-100. Both fractions were purified separately in the absence of any detergent using affinity chromatography with a Strep-Tactin XT column and a subsequent step of gel filtration using a Superdex200 Increase column (GE) in either 20 mM HEPES buffer with 20 mM NaCl at pH 7.4 or 20 mM Potassium Phosphate buffer with 20 mM NaCl at pH 7.4. SEC-MALS experiments were run in similar conditions injecting 5 mg/mL CLIC1 samples.

### NMR Spectroscopy

Purified ^15^N-labelled CLIC1 from both the membrane and soluble fractions were subjected to ^15^N-SoFast HMQC(27) or BEST ^15^N-Trosy experiments (28) at 30°C on a Bruker Avance3 spectrometer operating at a ^1^H frequency of 600 MHz or 800 MHz equipped with a TCI-P cryo-probe. High-field spectra were collected at the MRC Biomedical NMR Centre. Spectra were uniformly collected with 256 increments in the ^15^N dimension.

### Fluorescence Assays

Asolectin, a lipid extract from soybean, was solubilised in chloroform, dried under a stream of nitrogen and solubilised in HEPES or in phosphate buffer. 10 μM CLIC1 was incubated at 30 °C with 3 mM Asolectin lipids and was treated with 2 mM ZnCl_2_ or CaCl_2_ or left untreated, all in 50 mM HEPES 50 mM NaCl pH 7.4 buffer, or with H_2_O_2_ in phosphate buffer. Intrinsic protein fluorescence was recorded by excitation at 280 nm and emission was measured between wavelengths of 300 nm to 400 nm on a Varian Cary Eclipse fluorimeter, and between 400 to 500 nm with an excitation of 395 nm for the GFP-labelled samples. The samples were then spun at 208000 g for 30 minutes in an ultracentrifuge at 25 °C. Immediately after centrifugation, the soluble fraction was separated from the membrane pellet and the pellet was resuspended to similar volume as the supernatant. Tryptophan fluorescence was then carried out with the same methodology as described above for both the soluble and membrane protein fractions. All fluorescence data was normalised with subtraction of any background buffer or lipids.

### Chloride Efflux Assays

CLIC1 chloride channel activity was assessed using the chloride selective electrode assay described previously (4). Unilamellar Asolectin vesicles were prepared at 50 mg/mL in 200 mM KCl, 50 mM HEPES (pH 7.4). CLIC1 protein at 11 μM final concentration was mixed with the vesicles, incubated during 5 minutes and then 1 mM Ca(OH)_2_ or 1 mM ZnSO_4_ was added to a 2.5 mL final volume mixture and incubated again for 10 minutes. The lipid mixture was then applied to a PD-10 desalting column previously equilibrated in 400 mM Sucrose, 50 mM HEPES (pH 7.4) and collected in 3.5 mL of the same buffer. 500 μL of the lipid mixture were then added to a cup with 4 mL of 400 mM Sucrose, 50 mM HEPES, 10 μM KCl (pH 7.4) and the free chloride concentration was continuously monitored. 60 seconds after the addition of the lipid mix, 10 μM Valinomycin in ethanol was added and 60 seconds later, 1% TRITON X-100 was also added to release the remaining intra-vesicular chloride.

### Fluorescence Microscopy

Giant unilamellar vesicle formation was carried out using a protocol adapted from (29, 30). An Asolectin lipid stock was prepared in 50 mM HEPES, 50 mM NaCl pH 7.4 buffer. 2 μl/cm^2^ of 1 mg/ml lipid mixed with 1 mM Nile red lipophilic stain (ACROS Organic) was applied to two ITO slides and dried under vacuum for 2 hours. 100 mM Sucrose, 1 mM HEPES pH 7.4 buffer was used to rehydrate the lipids in the described chamber. 10 Hz frequency sine waves at 1.5 V were applied to the chamber for 2 hours. Liposomes were recovered and diluted into 100 mM glucose, 1 mM HEPES, pH 7.2 buffer. For all four assays 90 nM CLIC1-GFP was incubated with the GUVs with either 0.5 mM ZnCl_2,_ 0.5 mM CaCl_2_, or were left untreated and incubation at room temperature for ten minutes followed. Microscopy for each assay was performed in an 8 well Lab-Tek Borosilicate Coverglass system (Nun) with a Zeiss LSM-880 confocal microscope using 488 nm and 594 nm lasers. All images were processed with Zen Black software.

HeLa cells were kindly provided by Chris Toseland laboratory, University of Kent. HeLas were maintained in DMEM media supplemented with 10% FBS and 1% Penicillin/Streptomycin at 37°C, 95% humidity and 5% CO_2_.

For immunofluorescence assays HeLa cells were seeded into 24 well plates onto sterile microscopy slides for next day treatment. 24 hours post seeding cells were washed with Tris-buffered saline (TBS), transferred to phosphate free media and treated with 5 mM CaCl_2_ or 10 μM ionomycin as required. Fixation with 4% formaldehyde for 15 minutes was carried out at two hours post treatment. The cells were then washed and permeabilised for 10 minutes with 0.1% Triton in TBS and washed twice with TBS to remove any detergent. The cells were then stained with CellMask Deep Red plasma membrane stain (Invitrogen) at 1.5X concentration for 15 minutes. Following a TBS wash step the cells were blocked at room temperature with 2% BSA for 1 hour. Primary incubation was carried out overnight at 4 °C with a 1:50 dilution of monoclonal mouse CLIC1 antibody (Santa Cruz Biotechnology, clone 356.1). After primary incubation, 3 wash steps were carried out, prior to 1 hour incubation with secondary antibody at a 1:1000 dilution (Alexa Fluor 488 donkey anti-mouse, Life Technologies). A further 3 washes followed, then nucleus staining with NucBlu Live Cell Stain (Invitrogen) for 20 minutes. The slides were washed a final time and mounted with ProLong Gold Antifade (Invitrogen). All microscopy slides were viewed with a Zeiss LSM-880 confocal microscope using 405 nm, 488 nm, 633 nm lasers. All images were processed with Zen Black and Zen Blue software.

The fluo4 experiment was carried out from the same HeLa stock. The cells were seeded into 96 well plates and 24 hours later were treated with identical CaCl_2_ or ionomycin concentrations to the immunofluorescence assay, to verify intracellular calcium levels. Fluo-4 Direct (Invitrogen) was added to the cells at 1X dilution according to manufacturers’ protocol and visualised using a LS620 Etaluma microscope at the same time point as CLIC1 assay cells were fixed. Contrast and brightness were adjusted equally for all images and pseudo colouring was applied for intensity reading, using ImageJ.

### Mass Photometry

10µL of the protein/nanodisc was applied to 10 µL buffer on a cover slip resulting in a final concertation of 100nM. The data was collected on a Refeyn OneMP (Refeyn Ltd, UK) mass photometry system. Movies were acquired for 60 seconds. The mass was calculated using a standard protein calibration curve.

## ACKNOWLEDGEMENTS

We thank Dr N. Fili and Dr. J. Rossman for help with confocal imaging, Dr C. Toseland for providing HeLa cells, and Dr G.S. Thompson and Dr. D.A.I. Mavridou for feedback on the manuscript. We thank Refeyn Ltd for enabling us to collect our mass photometry data. We acknowledge the use of the MRC Biomedical NMR Centre, which is supported by Cancer Research UK (FC001029), the UK Medical Research Council (FC001029), and the Wellcome Trust (FC001029), via the Francis Crick Institute. We acknowledge support from the Wellcome Trust Seed Award (207743/Z/17/Z).

## CONTRIBUTIONS

JLOR, LV and ACH designed experiments. LV performed NMR data acquisition, fluorescence assays and chloride efflux measurements. ACH performed fluorescence assays, GUV experiments and cell imaging. EMC collected the mass photometry experiments. DC provided assistance with cell imaging experiments. JLOR, LV and ACH prepared figures. JLOR supervised the project and prepared the manuscript. JLOR, LV, ACH and DC edited the manuscript. LV and ACH contributed equally to the work.

