## Supplemental Figures for "Membrane insertion of soluble CLIC1 into active chloride channels is triggered by specific divalent cations"

**Figure S1. CLIC1 membrane insertion in *E. coli* is not driven by oxidation.** Overlay of $^{15}$N Trosy HSQC spectra of soluble CLIC1 non-treated (red) and reduced with 5 mM DTT (blue).

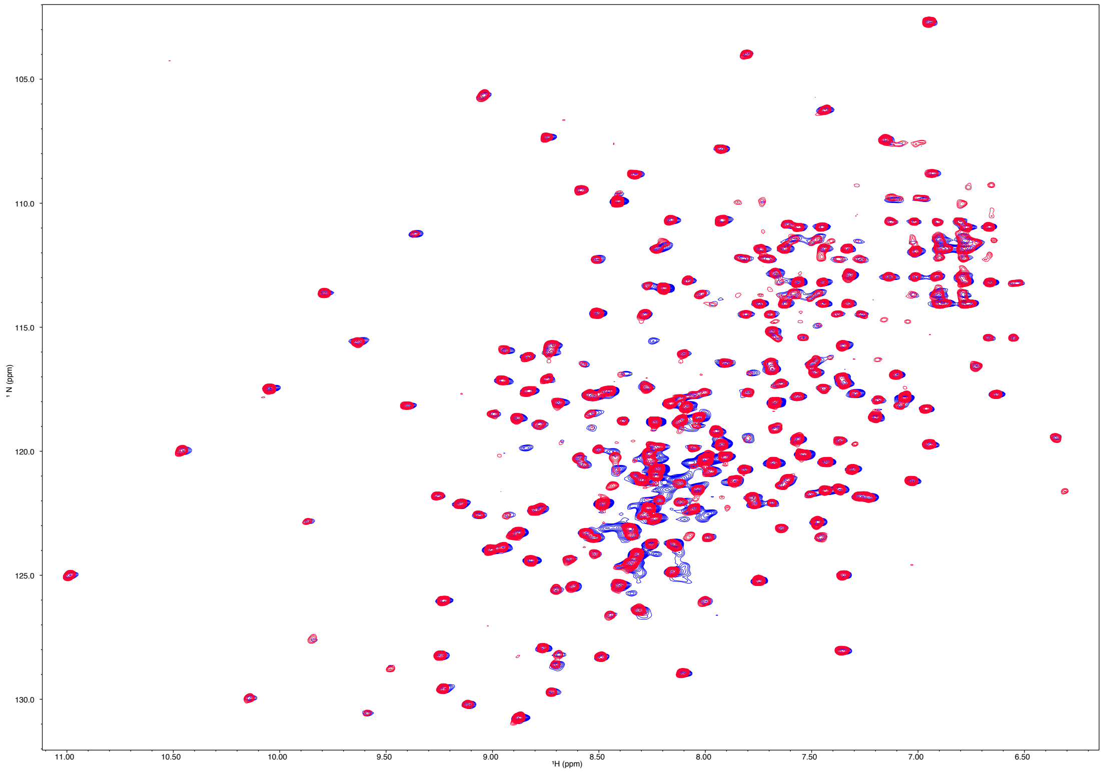

**Figure S2. CLIC1 association with lipids in the presence of divalent cations.** Native tryptophan fluorescence emission spectra of soluble CLIC1 (black) and the membrane fractions of CLIC1 incubated with asolectin in the presence of $Zn^{2+}$ (red).

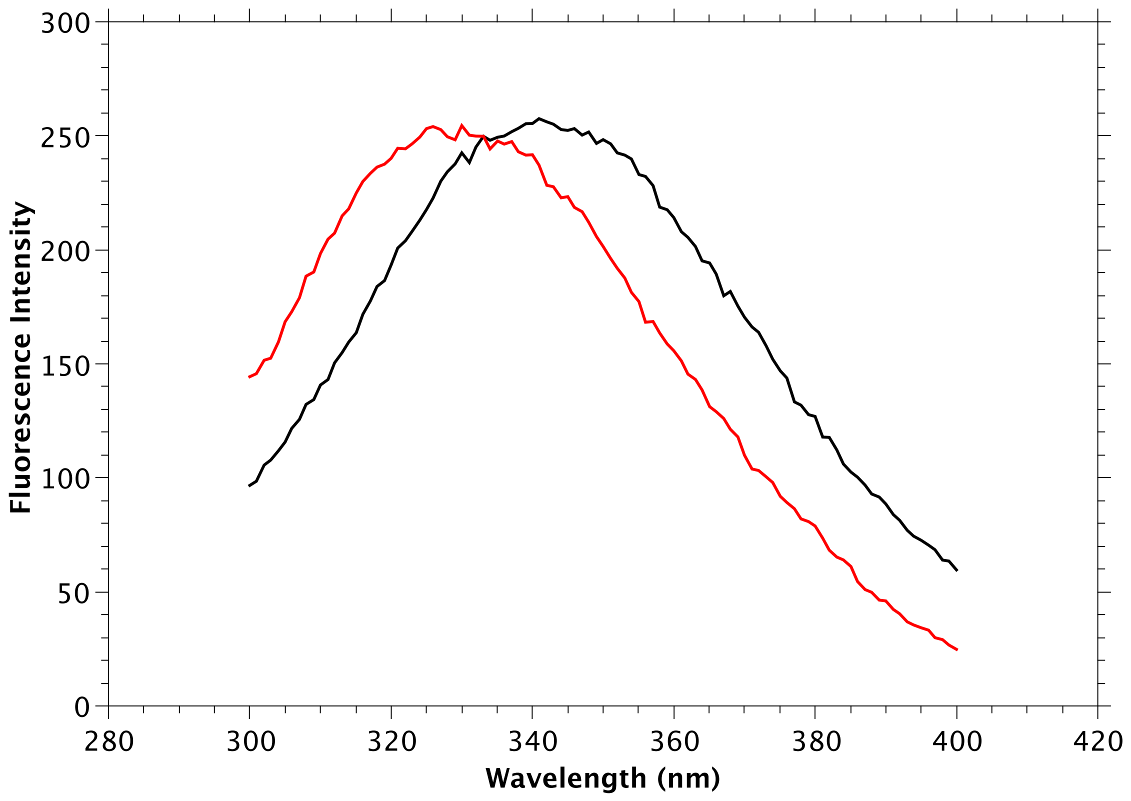

**Figure S3. Divalent cations trigger CLIC1 membrane insertion.** Widefield fluorescence microscopy images of Asolectin GUVs labelled with Nile red dye incubated with GFP-labelled CLIC1 in the presence of 500 μM of $Zn^{2+}$ (A,B) or $Ca^{2+}$ (C,D). The left panel shows images exciting Nile red, and the left panel shows images exciting GFP.

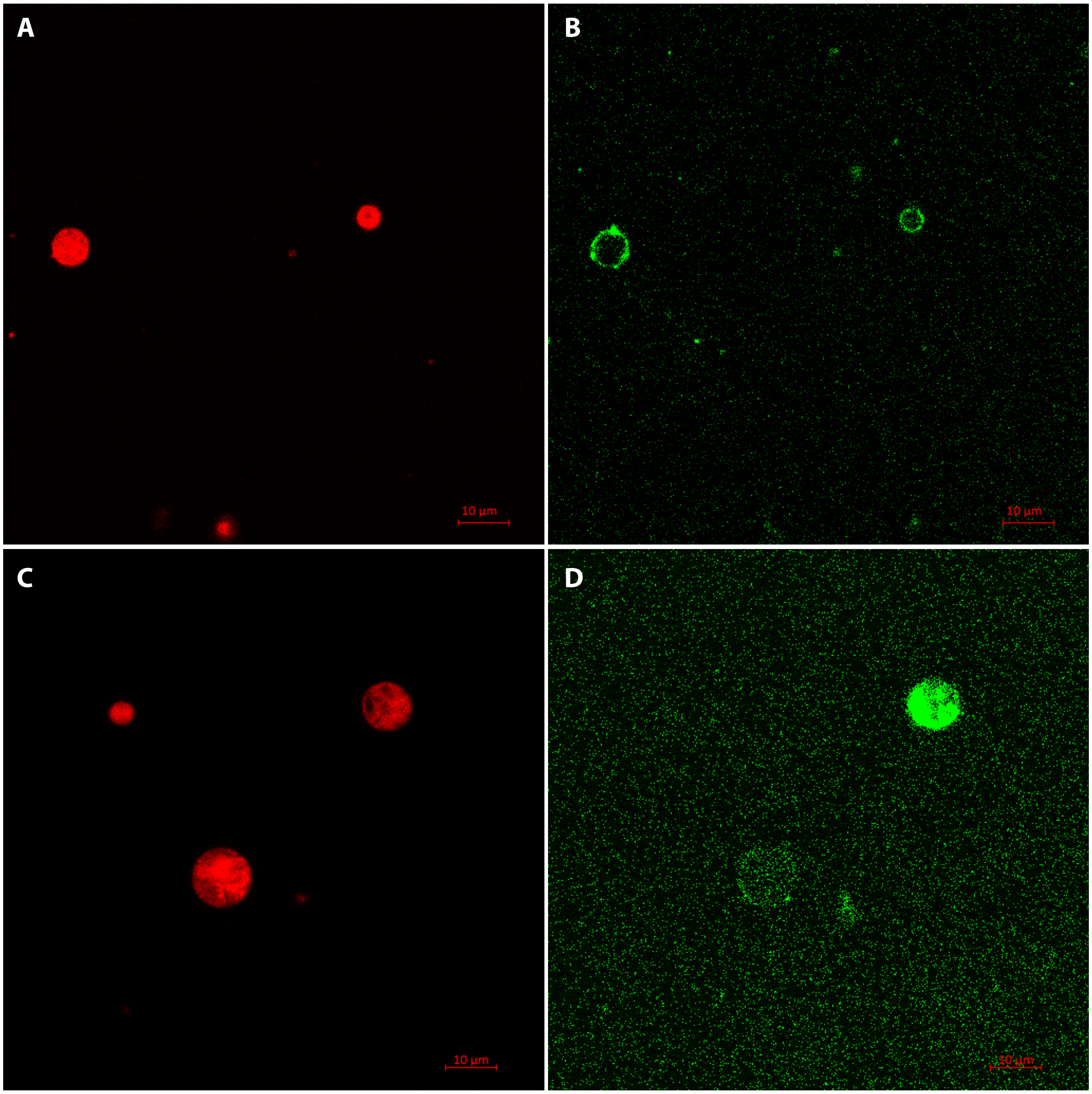

**Figure S4. Chloride efflux assay of CLIC1 channel activity**. CLIC1 chloride conductance monitored in Asolectin vesicles in presence of $Ca^{2+}$ (blue) or EDTA (red). A control experiment without the addition of CLIC1 is represented in black. The arrows indicate the addition of Valinomycin and Triton X-100.

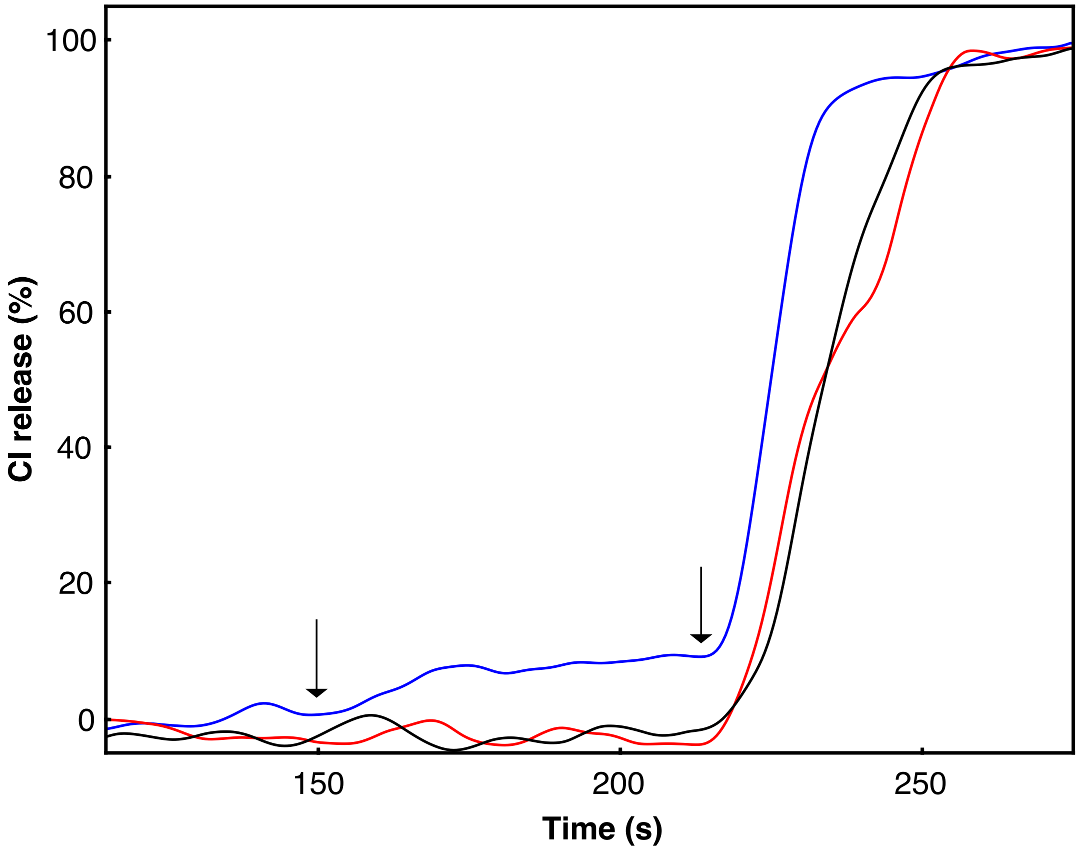

**Figure S5. Increasing the intracellular calcium levels in HeLa cells**. Intracellular calcium levels monitored following Fluo4 fluorescence (blue) in the absence (A) and presence of 5 mM $Ca^{2+}$ (B) and upon treatment of HeLa cells with 10 μM Ionomycin (C).

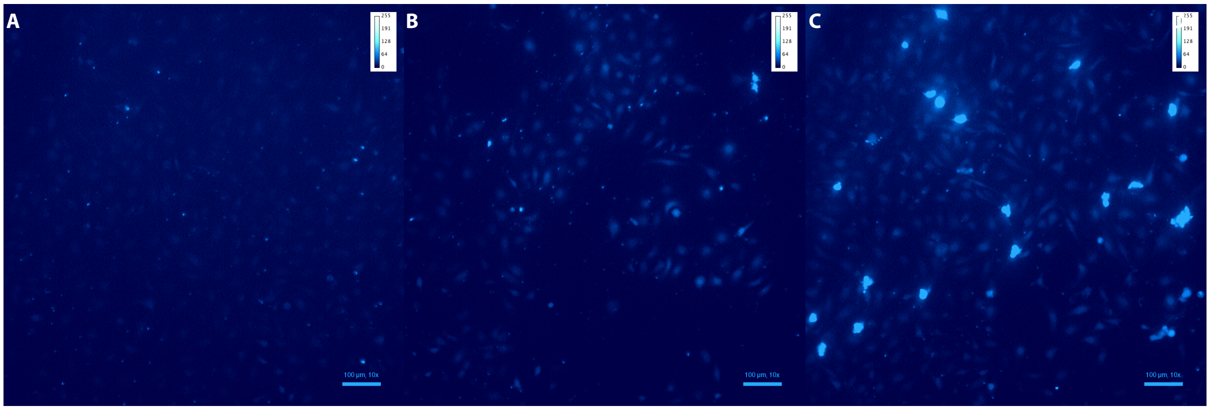

**Figure S6. $Ca^{2+}$ driven re-localisation of CLIC1 in HeLa cells**. Widefield fluorescence microscopy images indicating $Ca^{2+}$ driven re-localisation of CLIC1 in HeLa cells. Fluorescence microscopy images of HeLa cells stained with a CLIC1 antibody (green), a membrane marker (red) and DAPI (blue) in the absence (A-D) and presence (E-H) of 5 mM $Ca^{2+}$ in the media or upon treatment with 10µM Ionomycin (I-L). Images in D, H and L show a merge of the three different channels for the three conditions.

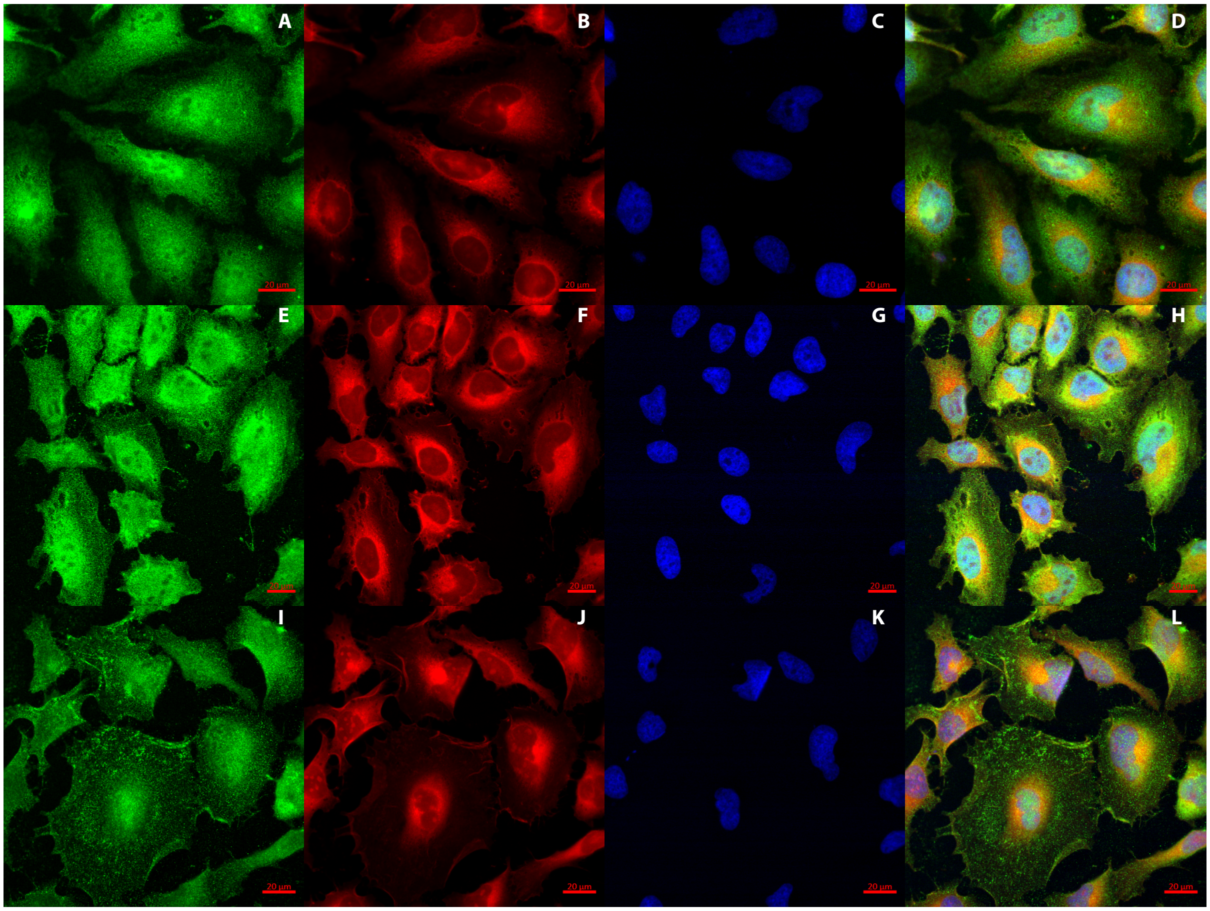

**Figure S7. CLIC1 oligomeric species are not dependent on cysteine oxidation.** SEC traces of non-reduced (black) and reduced (red) CLIC1.

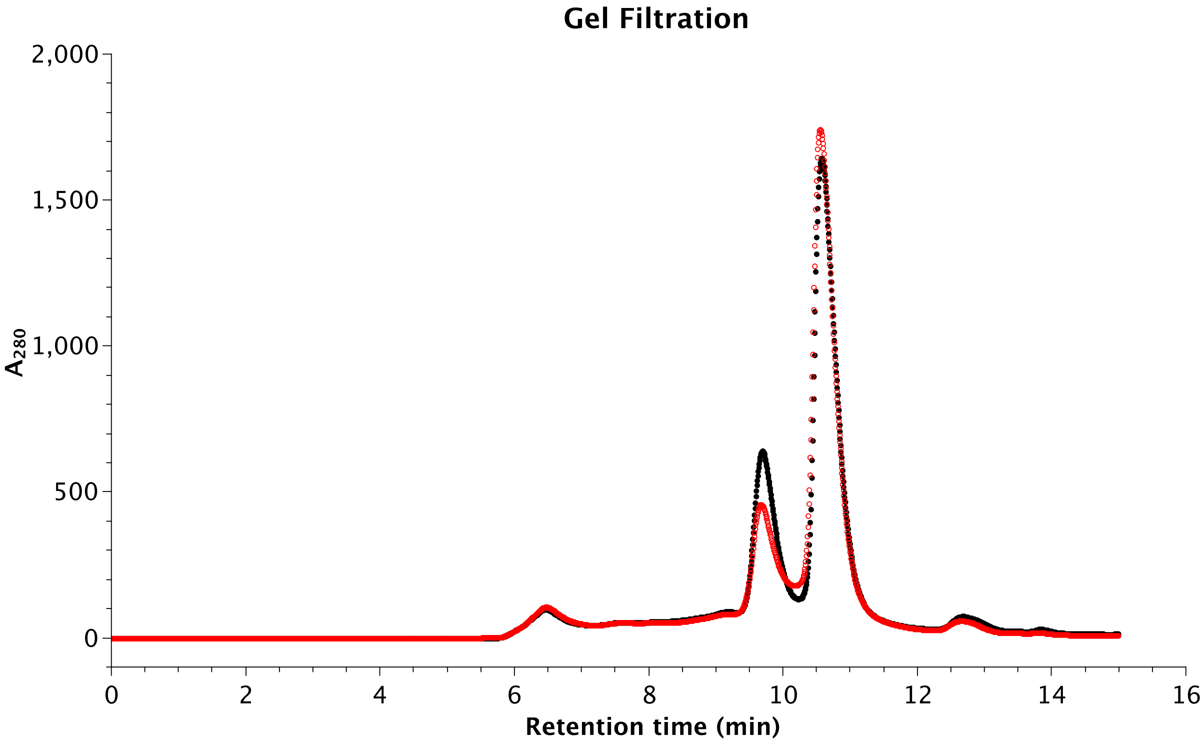
